# An investigation of social learning of tool-use in bottlenose dolphins (*Tursiops truncatus*) using a ball-up / ball-down task

**DOI:** 10.1101/2025.08.17.670717

**Authors:** Alenka Hribar, Annette Kilian, Claudio Tennie

## Abstract

Whether and how dolphins can engage in spontaneous, untrained social learning to solve novel tool problems via action- or result- (or other) social learning remains debated. In the present study we tested the spontaneous social learning abilities of six dolphins (all not trained to copy) on two tool-using tasks (what we call the ball-up / ball-down task) - using a dolphin demonstrator and a human demonstrator. Regardless of task type and demonstrator type none of the tested dolphins reproduced the demonstrated tool solutions. We experienced several issues regarding our test apparatus, and so these negative results may be due to apparatus failures. However, these findings may also fully or partially indicate that untrained dolphins are not generalized, spontaneous social learners across information types, especially regarding the acquisition of tool solutions in puzzle tasks. More studies are required to precisely determine the capacity for spontaneous tool solution copying in dolphins.

## Introduction

All cultures are (directly and/or indirectly) social enterprises – culture is dependent on the presence of effects of some type(s) of social learning. Social learning refers to the acquisition of information from *or via* another individual or its behavioral products (Heyes 1994). Social learning can take place via multiple social learning types, which are not well understood on a neurological level but are typically differentiated based on the nature of the information acquired (see reviews, e.g. by Whiten and Ham 1992). A class of information that involves information about the biomechanical actions involved in a behavior and/or the artifact or artefact movements resulting from the behavior is *know-how* (where the know-how can be sequentially and/or hierarchically organized, too; Tennie et al., 2017). Social learning of know-how may trigger the release and/or development of latent know-how in observers, or it may lead to copies of know-how that the observers would (absent the observation) have been very unlikely to perform (Tennie et al. 2017). Subtypes of social learning of know-how are imitation, where observers act similar to demonstrators following their observation (e.g. Zentall, 2006). Another subtype is emulation, which can can refer to the reproduction of the outcomes (or, more psychologically oriented: the goals) of a behavior (Tomasello 1990; see Huang & Charman, 2005 for an overview of emulation’s subdivisions), but using one’s owns means. Emulation learning itself can take place via goal-emulation (adopting an inferred goal) and/or affordance learning (learning about the physical properties of the environment and relations among objects) and/or object movement reenactment (replicating what objects did, i.e. how they moved; reviewed by Hopper, 2010). In all these cases, the distinction between triggering and copying can be upheld as described (e.g. the subtype imitation can lead to a trigger or to a copy of action patterns).

Humans can copy know-how even if this know-how is beyond potential individual reach and such *know-how copying* has been deemed a necessary enabling condition in human cultural evolution (Boyd & Richerson, 1996; Tomasello, 1999; Tennie et al. 2017). The existence of know-how copying abilities in non-human animals, however, is a much more debated topic. Claims of know-how copying abilities are particularly common for great apes and cetaceans (especially odontocets, Kuczaj II & Yeater, 2006). Chimpanzees (*Pan troglodytes*) for example have been suggested to *have to acquire* (thus, a claim for a need for copying) some of their cultural repertoires (patterns of behavioral variation within a species that are not directly correlated with genetic or environmental differences between populations; see e.g. for some such claims inside Whiten et al. 1999, 2001); additional claims of know-how copying abilities of chimpanzees are common in the primatological literature (e.g. reviewed in Whiten et al. 2009). However, other studies have suggested, based on experimental data, that great apes do not spontaneously develop marked know-how copying abilities outside of human influence (Tomasello & Call, 1997; Tennie et al. 2012; Neadle et al. 2020; Tomasello et al 1993b).

Multiple studies attribute social learning, and also copying, of know-how – both via imitation and emulation – to dolphins, particularly to bottlenose dolphins (*Tursiops truncatus* and *Tursiops sp.*; reviewed by Kuczaj II and Yeater 2006; Herman 2002; Kuczaj et al. 2012). Potential evidence comes from four sources: first, there are studies looking into copying vocal know-how. Second, there are studies on synchrony in dolphin behaviour, which is often linked to action know-how copying skills; third, several anecdotal reports have been linked to type of know-how copying; fourth, experimental studies have reported results congruent with action know-how copying in dolphins. Fifth, other experimental studies claim for results copying in tool use tasks. Sixth, there are studies on social learning of tool use in the wild. We will discuss these sources in turn. But first, we note that the claim that dolphins have some social learning of some type(s) of information is not an extraordinary claim. Social learning in animals is extremely widespread, and it would be surprising to find that dolphins are incapable of any type. Indeed, as we will see, the below provides good evidence for some types of social learning in bottlenose dolphins.

### Vocal behavior

Evidence exist for social learning of vocal behavior, such as so-called signature whistles (evidence in bottlenose dolphins: King et al. 2013). Particularly strong and controlled evidence for the spontaneous social learning of vocal know-how in dolphins stems from a study on wild Atlantic spotted dolphins (*Stenella frontalis*, Herzing et al. 2024). Several of these wild dolphins spontaneously (albeit partially) copied several computer-generated sounds (CGS) – including even the “start” and “stop” tones that preceded and followed the CGSs that these dolphins were exposed to. These computer generated sounds (and even the start and stop tones) were novel to there dolphins and would highly unlikely have been used or developed by these dolphins lacking the demonstrations (as the authors state these sounds “were designed to be outside the dolphin’s natural repertoire” and this was tested against baseline data). The fact that these wild dolphins (though note that they had years of human contact) nevertheless copied these sounds – absent human training and (possibly) absent any human enculturation - renders the results of this study a relatively clear case of spontaneous vocal know-how copying in dolphins. Thus, this is not only clear evidence of social learning in dolphins, but also clear evidence for know-how copying. However, the ability to copy in one domain (here the vocal domain) need not entail similar abilities in other domains. We will thus turn to these other domains.

### Synchronous behavior

Adult dolphins spend a considerable amount of time in synchrony with other dolphins (Connor *et al*. 2000; Connor *et al*. 2006; Fellner *et al*. 2006; Perelberg & Schuster, 2008). Why dolphins synchronize is not altogether clear but previous studies have suggested that the coordinated breathing of dolphins represents an act of cooperation (Perelberg & Schuster, 2008), and that male bottlenose dolphins might surface synchronously as an alliance signal (Connor *et al*. 2006). In some ways, even the fact that dolphins can form social groups is in itself evidence for some social learning: the very location of dolphins in time and space is clearly influenced by other dolphins – thus satisfying the minimal definition of social learning given above. Moreover, however, dolphins frequently exhibit spontaneous synchronous group behavior while traveling, foraging, playing, resting and displaying (Connor *et al*. 2000; Connor *et al*. 2006; Fellner *et al*. 2006; Miles & Herzing, 2003). Furthermore, during the first three months of a newborn’s life, a mother spends 80% of her time in synchrony with the newborn (Fellner *et al*. 2006; Mann & Smuts, 1999; Miles & Herzing, 2003). As social behavior and synchronous behavior appear in wild dolphins, dolphins clearly perform synchronous behavior spontaneously – in the sense of not requiring human influences to develop this aspect of their behavior. It has even been suggested that imitation (or more neutrally: social learning regarding actions) in dolphins might spontaneously develop first from passive and later from active maintenance of such synchrony (Fellner *et al*. 2006).

Synchronous behavior in dolphins has also been tested in captivity, though after training and human interference. Two captive bottlenose dolphins were human-trained to perform a particular behavior in tandem (i.e., in synchrony) upon receiving a signal from their trainer (Braslau-Schneck, 1994; as cited in Herman, 2002). Additionally, the same dolphins were later signaled to perform an unspecified behavior in synchrony. In both cases, the dolphins executed the behaviors in almost perfect synchrony (after both dolphins had first spent several seconds swimming together underwater), with one dolphin always performing slightly ahead of the other dolphin (Braslau-Schneck, 1994; as cited in Herman, 2002).

How exactly captive and wild dolphins achieve synchronization, remains debated. One possibility could be that a slightly delayed dolphin copies (or gets triggered by) another dolphin’s actions or muscle innervations. However, it is not clear whether synchronous behavior represents a case of this *per se*, as social learning about others’ actions might not be a necessary requisite in order to synchronize – note also that not all muscle innervations are easy to observe or infer (all this in a potentially moving medium of water). Synchronous behaviors may instead occur as a consequence of the lagging dolphin closely following (in parallel) the direction of movement (or a parallel position’s match) of another dolphin without necessarily copying the leading dolphin’s actions or muscle innervations (that is, instead of copying or triggering action know-how, dolphin synchrony may be due to social learning of “know-*where*” another dolphin is across time (know-when) perhaps in relation to oneself; compare Bandini et al. 2020, Tennie et al. 2020). Overall, the available data on synchrony therefore does not unquestionably demonstrate a role of action / muscle social learning (be it triggering or copying) in the performance of synchronous behavior and in particular it does not demonstrate that observers copied bodily actions that they themselves could not have developed on their own absent demonstrations.

### Anecdotal observations

Anecdotal evidence for action social learning in dolphins includes instances of dolphins (bottlenose dolphins and one false killer whale, *Pseudorca crassidens*) claimed to reproduce actions performed by other dolphins (Brown *et al*. 1966; Caldwell *et al*. 1965; Pace, 2000; Tayler & Saayman, 1973) as well as reproducing actions performed by individuals of other species (humans, Cape fur seal, fish, penguins, skates and loggerhead turtles, see Tayler & Saayman, 1973; Kuczaj et al. 2012; for a review see Herman, 1980). Bossley et al. (2018) reported a case in which a wild dolphin was temporarily housed with captive, trained dolphins. Once released, the dolphin started performing a behavior (tail walking) that was present (human-trained) in the captive population but was never observed before in the wild. Furthermore, years later, another wild dolphin started to perform tail walking behavior. Although these observations provide evidence for the existence of social learning in dolphins (especially the second wild tail walker, who presumably never had much human contact), they do not provide conclusive evidence of the social learning of actions (though they are consistent with this possibility). Namely, it is again unclear what kind of information these two dolphins acquired from or via the demonstrators. Instead of copying the actions, perhaps these dolphins learnt more generally that their bodies could be maintained outside of the water (“know-where” information; or maybe the learning related to the goal of moving the body out of the water) and they then subsequently reinnovated the behavioral form itself (tail walking) on their own – e.g. by trial and error learning.

Whilst anecdotal evidence can be helpful insofar as it can guide the development of experimental studies or more systematic observations, anecdotes can hardly be regarded as a research strategy on their own (Maestripieri & Whitham, 2001) and systematic studies are needed to test the hypotheses stemming from anecdotes. We therefore note that these anecdotes are of interest and consistent with the idea of action social learning (and perhaps even action copying), but they alone cannot prove this as too many alternative possibilities remain. Yet, they once again show the presence of spontaneous social learning in wild dolphins.

### Experimental studies on actions

The fourth source of information about dolphin know-how social learning abilities results from experimental studies on action social learning. As alluded to already, experimental studies on captive dolphins often have the limitation that the individuals may not be ecologically representative of their wild counterparts because they may be human-enculturated (henceforth: enculturated) and/or human-trained. Data obtained from such studies may not capture what dolphins may do spontaneously, absent human interference. Enculturation refers to animals that have been reared in a human cultural environment with wide exposure to human artefacts and/or social/communicative interactions (Furlong et al., 2008). Enculturation has been suggested to be able to principally induce cognitive abilities that are not present and would not develop otherwise in wild, untrained conspecifics - such as the ability to readily *copy* actions in great apes (Henrich & Tennie, 2017; Tomasello & Call, 2004; Tomasello et al. 1993b).

Two studies (Herman et al. 1989; as cited in Herman, 2002; Xitco, 1988) used the so called “Do as I do” paradigm to investigate copying abilities in bottlenose dolphins. By necessity, in the “Do as I do” paradigm, a subject is first *human-trained* (by shaping, for example) to reproduce some demonstrated actions on command (the command often being the words “Do this”, or an equivalent gestural signal). The subject is then tested on transfer actions (which often also include the originally trained actions – but really should be novel, and ideally, actions that would not otherwise occur). The study by Xitco (1988) involved two bottlenose dolphins (Phoenix and Ake) who acted as demonstrator and as observer for each other (Xitco, 1988). The transfer actions were either from the preexisting repertoire of both dolphins (familiar actions - thus not fully improbable “transfer” actions) or they were previously taught to only one dolphin - the demonstrator - and so were considered novel for the *other* dolphin - the observer (*i.e.,* these were *assumed* to be “novel” actions, but see Byrne and Tanner, (2006) for a general critique of such an approach). Familiar actions were reproduced. Phoenix successfully reproduced 7 of 12 of the familiar actions and Ake successfully reproduced 6 of the 12 familiar actions. Phoenix also reproduced 2 of the 3 novel actions (both on the second trial) and Ake reproduced 1 of 3 of the novel actions (on the third trial).

A later study using the Do as I Do paradigm on the same two dolphins (Herman et al. 1989; as cited in Herman, 2002) also compared the repetition rate of actions depending on whether the demonstrator was a dolphin or a human. The authors found that there was no significant difference in action repetition by these two trained dolphins depending on whether the demonstrator was a dolphin (62% correct) or a human (58% correct). Although these rates of repetition might seem high, it is important to take into consideration that both dolphins tested by Xitco (1988) and Herman et al (1989) had also experienced years of language-training where they were required to produce various actions on command and that they had ample experience in performing actions in tandem on command. Given that these dolphins received approximately eight to ten hours a day of human exposure (including training and cognitive experiments), these individuals should be considered not merely human-trained but also human-enculturated (Herman, 2002).

While the aforementioned studies show some potential under special circumstances (human interference) for action *copying* in bottlenose dolphins, their ecological validity is clearly compromised both by their degree of training and enculturation of the tested subjects. It is possible, if not likely, that the detected action copying abilities might have been *induced* in the two tested dolphins through extensive training *and/or* enculturation.

Bauer and Johnson (1994) set out to replicate Xitco (1988) study in order to test the copying abilities of bottlenose dolphins who had not had years of prior specialized training. The subjects in Bauer and Johnson’s study (two bottlenose dolphins named Toby and Bob) were tested using the same experimental procedure as Xitco (1988). Although Toby and Bob learned most of the training actions and almost all of the transfer actions a few months prior to the study in the Do as I Do paradigm, the two dolphins were described to have had worse reproduction rates than those dolphins tested by Xitco (1988) when asked to replicate familiar actions. Moreover, when asked to replicate *novel* actions, neither of the two dolphins replicated any of the actions correctly (Bauer and Johnson, 1994). Based on these results, Bauer and Johnson concluded (in the terminology of the current paper) that their two dolphins lacked action *copying* skills. Consequently, it is possible that only dolphins that have experienced enculturation and/or other, specialized training can copy – mere training (as in Do As I Do training) may not suffice. Of course, potentially many other differences may explain these discrepant results. Overall, it therefore remains an open question whether dolphins can easily and spontaneously (learn to) *copy* know-how (here: action know-how) – though the study by Bauer and Johnson (1994) suggests that they do not. What is remarkable is that even following some human-*training*-to-copy, dolphins can still fail to copy novel actions (Bauer and Johnson 1994).

### Experimental studies on environmental results

Regarding the question of whether dolphins can copy environmental results (a variant of emulation; see above), we are aware of only one such study that contained a claim of such abilities (sensu Kuczaj & Walker, 2006). Captive bottlenose dolphins were reported to spontaneously copy results achieved by human demonstrators in a problem-solving task. In particular, in one of the experimental conditions called the “multiple-weight task”, dolphins had to pick up and later drop four weights into a container in order for a fish reward to be released, which they allegedly learned by observing human divers’ results-demonstrations (sensu Kuczaj & Walker, 2006). Given that the target of the task was to place the weights (know-what; Tennie et al. 2020) in a precise location (know-where; Tennie et al. 2020) rather than to investigate if the dolphins could perform the actions involved in solving the task (which they were already trained to do, Kuczaj *et al*. 1998), reproduction of the solution demonstrated by the divers could evidence emulative abilities in dolphins in the sense of know-how, or else the social learning of a combination of know-what and know-where (see Tennie et al. 2020). In addition, as the task was not novel to the dolphins, it is not possible to rule out carryover effects from previous experiments – meaning the novel, unlikely know-how transmission can be ruled out (not a test for results *copying*).

In another experimental condition of the same study (sensu Kuczaj & Walker, 2006) called the “2-Step Time-Limited task”, the same dolphins that participated in the previous condition had to now use two tools (a weight and a stick) in quick succession (15 s) in order to get to a fish reward out of a puzzle box (sensu Kuczaj & Walker, 2006). Again, the authors mention (but do not elaborate) that dolphins learned to use these tools by observing a human demonstrator. Later during the experiment, the stick-tool was placed further away from the tool site and the dolphins had to bring it closer to the apparatus before they used the weight in order to also be able to use the stick within the limited time frame. None of the dolphins spontaneously brought the stick close to the puzzle before using the weight box. However, the authors report that after one of the dolphins saw a human model bringing the stick to the puzzle box before using the weight, one dolphin “quickly began to do so himself” (sensu Kuczaj & Walker, 2006). What, if and how the dolphins actually learned to solve the task is again unclear – especially given that the amount of training they had received is not described in detail. Know-how copying may be a possibility, in the sense of results copying. But other types of social learning may have acted instead or in addition (such as social learning of know-what and know-where and know-when). Overall, then, none of the studies to date conclusively show that dolphins spontaneously copy either novel actions or novel results (*know-how copying*) – but tool use tasks (such as the weight or weight and stick task) seem to hold some promise. In general, social learning can take place across several behavioral domains but in the present study, and from here on, we shall focus on the technological (here: tool) domain.

### Tool use in wild dolphins

Some individuals in populations of bottlenose dolphins have been described to use tools in foraging contexts. One tool use behavior described in bottlenose dolphins, sponging, was reported in Shark Bay (Australia) (Krützen *et al*. 2005; Mann *et al*. 2008; but see also Parra, 2007 for observations on one sponge carrying Indo-Pacific humpback dolphin). Sponging consists in the use of conical sponges likely as “gloves” for the rostrum while foraging for buried prey in the sand. Sponging behavior is highly biased towards females and recent studies have shown that genetic and environmental factors do not account on their own for the pattern of distribution of this behavior within matrilines, suggesting that (vertical) social learning of some type(s) is present in sponging behavior (Wild et al. 2019; Krützen et al. 2005). This leaves open the precise social learning type(s) that may be involved. A second tool use behavior described in wild bottlenose dolphins is shelling (Allen et al. 2011). During shelling, dolphins guide prey into empty gastropod shells or directly feed on prey hiding in such shells by carrying the shells to the surface, emptying the water and shaking the shells (Allen et al. 2011). Recent analysis again incorporating both genetic and environmental data have shown that contrary to sponging, shelling, too provides evidence for some social learning. In shelling, the social learning was found not to be vertical but rather to horizontal or oblique, as in social learning between peers (Wild et al. 2020). Again, this leaves open the social learning type(s) that may be involved.

There are more types of tool use in dolphins than the two described (e.g. among them so-called mud-ring feeding (Torres & Read 2009, Engleby & Powell 2019) and also social tool use in cooperation settings – even across species (Simões-Lopes et al. 1998). Unfortunately, none of the observations of tool use in wild dolphins allow for easy assessing which social learning types accompany, underlie or perhaps have to underlie (the latter a claim for copying) the acquisition of these tool-based foraging techniques. Given that the action social learning capacity of dolphins has been mainly addressed using demonstrations of behaviors already within the dolphins’ repertoire of bodily movements, and that the tool use tests performed in captivity are inconclusive (see above), we may say that “Social learning of relatively novel techniques of food-handling […] has yet to be demonstrated.” (Whiten & van Schaik, 2007, pg. 615) using “ well-controlled experiments” (Mann *et al*. 2007; Sargeant *et al*. 2005) – in the sense of the techniques themselves being socially transmitted.

A firm answer to what type(s) of social learning may or may not be spontaneously present in dolphin tool behavior – and especially which type(s) may be necessarily required – is far off at the present time. However, to provide better and firmer answers, more studies are required, that systematically vary information types in demonstrations to determine which types of information are and which are not (or to a lesser degree) socially learned by dolphins. As one additional important question related to the presence of absence of know-how copying types (as defined above), such studies should also ideally test the baseline competencies in motivated subjects. In all cases, to attain ecologically and phylogenetically meaningful answers, tests should be conducted that test for spontaneous or spontaneously developing abilities – that is, untrained and ideally unenculturated subjects should be tested.

The present study was designed to test for the spontaneous social learning abilities of two types of tool-use behaviors by bottlenose dolphins (*Tursiops truncatus*). We conducted these two studies with bottlenose dolphins who had mild levels of enculturation (at least when compared to some of the earlier tested dolphins) and who had also not been trained on any of the actions or results needed to solve the target tasks. We adapted parts of the experimental design of the “multiple-weight task” used by Gory and Kuczaj (sensu Kuczaj & Walker, 2006) to simplify the task given the lack of preliminary training in the tested dolphins. The result we label the ball-up / ball-down task. Subjects were required to release only one “weight” (here: a ball; this being the know-what) - instead of four - in order to obtain a fish reward. In the first study (the ball-up variant of the task), the dolphins had at their disposal air-filled balls (know-what 1) that could be used to retrieve the fish by releasing them at the lower end of the tube (*i.e.*, the ball rising in the tube would displace the fish out the other end; know-where 1). In the second study (the ball-down variant), the dolphins were given negatively buoyant balls (know-what 2) that could be placed in the top of the tube (know-where 2) thereby having them sink through the tube and displacing the fish at the bottom. In the first task the demonstrator was a dolphin (who demonstrated target results as well as target actions), and in the second task the demonstrator was a human (who modeled target results but not target actions, since she used her hands). We report on these two studies now, although we note that we experienced some methodological issues with our setting, which prevent conclusive claims being drawn.

## Study 1 – Ball up (dolphin demonstrations)

### Materials and Methods

#### Subjects

Subjects were six bottlenose dolphins (*Tursiops truncatus*) housed in the dolphinarium at the Tiergarten Nürnberg (Germany) (details Table 1). The dolphins were kept together with six California sea lions (*Zalophus californianus*) in three indoor tanks connected to each other through short passages. The experimental sessions took place in a circular tank 12m in diameter and 4m in depth (from now on called the “testing tank”). Its adjoining tank (25m x 11m, 4.7m deep) was used for public performances and training. The third tank (9m x 5m, 2.5m deep) was on the other side of the “performance” tank. The dolphins were fed five to six times a day and were never food deprived during the study. At the time, the dolphins participated in four to five trainings or public performances a day, depending on the season. One dolphin, Noah, had previously participated in two studies on numerical competence (Kilian *et al*. 2003; Kilian *et al*. 2005).

**Table 1.** Dolphins that participated in Study 1 and 2.

| Dolphin | Age (years) | Born where | In Nürnberg since |
| --- | --- | --- | --- |
| Sunny | 7 | Soltau | September 2005 |
| Naomi | 8 | Nürnberg | birth |
| Noah | 13 | Nürnberg | birth |
| Jenny | 21 | wild | 1991 |
| Eva | 37 | wild | 1979 |
| Moby | 46 | wild | 1971 |

Note. Ages for wild born dolphins are approximate.

#### Apparatus

The testing apparatus was a clear hollow polycarbonate transparent tube (90cm long and 23cm in diameter) open on both ends. This apparatus was baited with a fish inside and attached vertically to a metal frame at the edge of the testing tank. The fish-bait was attached in the middle of the tube with a white plastic lacing cord. The shallowest opening of the tube was fixed at a depth of 30cm from the water surface (see fig. 1).

**Figures. 1a and 1b.**
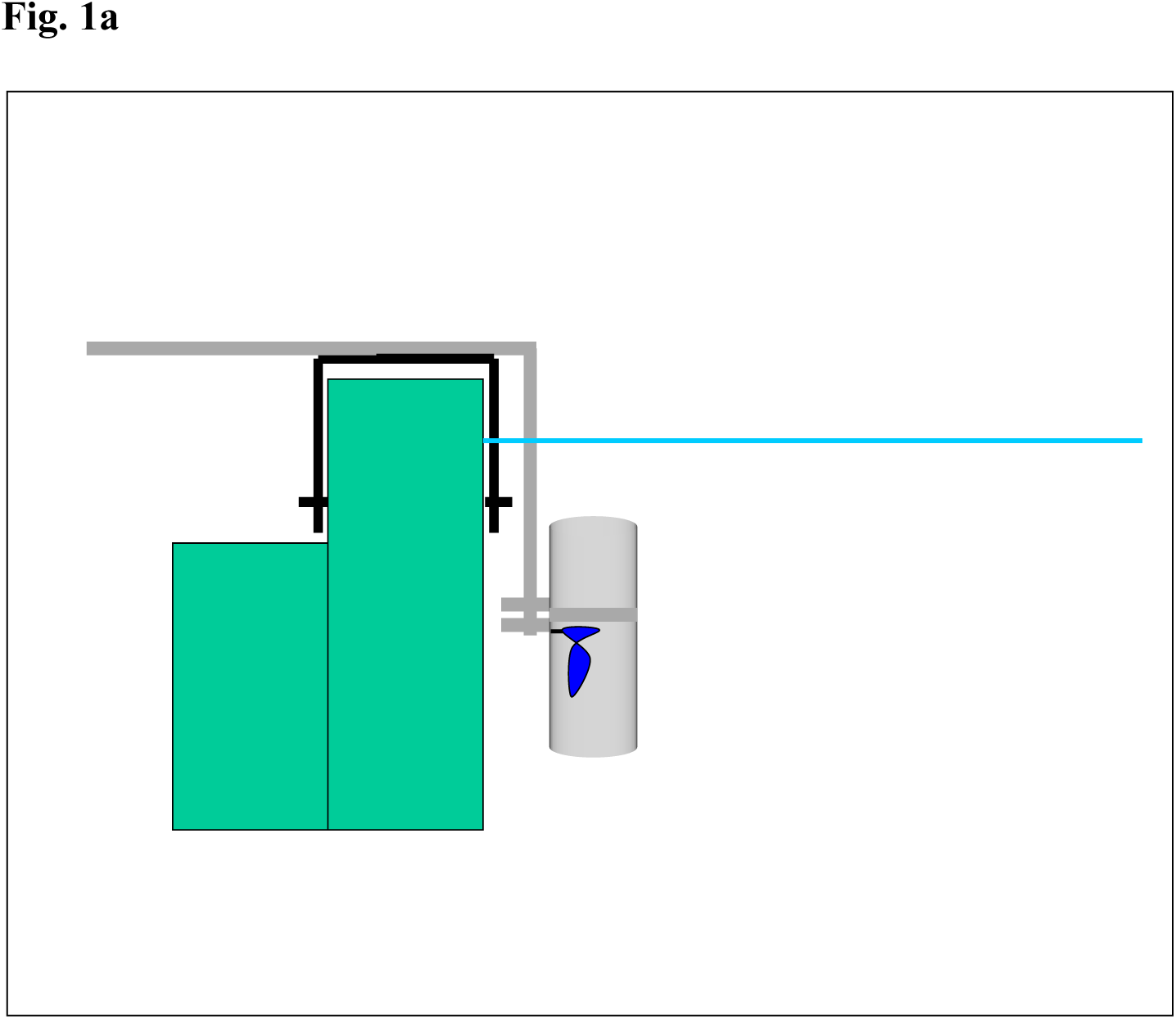
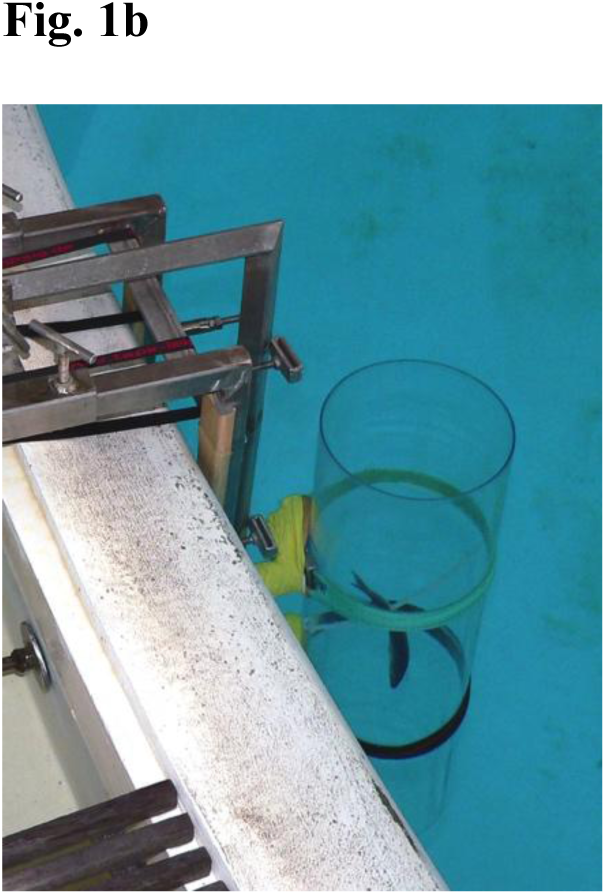
Apparatus. To get the fish-reward in Study 1 (Ball up) the subjects had to position the air-filled ball (know-what 1) under the tube’s lower opening (know-where 1) and let it go. The ball then rose to the water surface and untangled the fish from the lacing cord on the way up. In Study 2 (Ball down) they had to bring the heavy ball (know-what 2) over the tube’s upper opening (know-where 2) and let it go. The heavy ball then sank through the tube to the tank’s floor and again untangled the fish and brought it with it when it came down. *Note* Figure 1a is not to scale.

The “tools” provided to the dolphins were small air-filled basketballs (know-what 1; 15cm in diameter). The dolphins knew these items, as they were often used during trainings/performances by throwing them out of the pool to the trainer or balancing them on their rostrum. The dolphins also played with such balls extensively outside of training sessions by carrying them in their mouth, throwing them above the water or out of the tank or swimming under water with them.

In the demonstration phase for Group 2 (see Procedure section), a target was fixed to the tube. A target is a regular training tool used to shape an animal’s behavior. In this case the target was a stick (130cm) with a tennis ball attached to the end vertically fixed to the outside of the tube and taped at both ends so that the tennis ball rested just below the bottom lip of the tube.

Before the study started, we verified whether the dolphins could perceive the (transparent) tube by throwing a similar tube (smaller in diameter and length but of the same material) into the testing tank. Once the tube reached the bottom of the tank, the trainer instructed the dolphins to “bring the object back”. Given that the tube was immediately retrieved by the dolphins, we assumed that the tube was visible and could be perceived by the dolphins. The dolphins could also clearly perceive the fish in the tube, as was apparent from the fact that when they were let for the first time into the testing tank with the baited apparatus present (baseline phase), they swam directly towards the fish and tried to extract it.

#### Testing Procedure

One above-water camera was positioned at the edge of the pool some meters away from the apparatus in order to film both the apparatus and the area around it (approximately 2 meters in diameter).

Study 1 consisted of four phases: 1.) baseline, 2.) training of a dolphin demonstrator, 3.) target control, and 4.) trials with demonstration. The baseline phase was conducted with all the dolphins together. In the demonstrator training phase, the dolphin acting as demonstrator was separated from her group. In the target control and the trials with demonstrations the dolphins were divided into two groups. Group 1 comprised of Moby, Eva and Sunny and Group 2 of Noah and Naomi. We tested the dolphins in groups instead of individually, because the dolphins were not used to being separated from the group (except for medical reasons).

- *Baseline.* The baseline phase consisted of five 15 minutes sessions and was designed to familiarize the subjects with the experimental apparatus and to document the actions and choices displayed spontaneously by the subjects (including spontaneous occurrences of the use of the balls as tools). At the beginning of each baseline session, the baited apparatus and six balls were placed in the water before the dolphins were let into the testing tank. This baseline acted as a test for whether the dolphins required copying to solve the task.
- *Training of the dolphin demonstrator*. Jenny was used as the dolphin demonstrator, and so she was trained. For the time of training she was separated from the rest of the group. We used targeting to train Jenny to use the ball as a tool to get the fish out of the tube by letting the ball float upwards from the lower tube opening. Targeting is a shaping technique where an animal is taught to touch some part of its body with an object, called the target (Ramirez, 1999). The target was used to guide Jenny toward the tube’s lower opening. At the beginning of the training phase, Jenny was given a ball and a new hand signal, and then she was guided by the target to the lower hole of the tube. The training phase consisted of 10 sessions, each with 10 to 16 trials.
- *Target control*. Given that Jenny needed the guidance of a target in order to be a successful demonstrator, we added one control session to control for possible effects of the target on the dolphins’ behavior toward the tube (as compared to the baseline). The procedure was exactly the same as in the baseline sessions and each session (1 per group) lasted 10 minutes. For Group 1 the experimenter held the target in the exact same position as she held it for Jenny during the demonstrator training phase but for Group 2 the target was fixed to the tube.
- *Trials with demonstrations*. Jenny served as the demonstrator for both groups during four sessions in each group. Each demonstration started by giving Jenny a ball and the hand signal to start the target behavior (bring ball to lower opening of the tube and release the ball). During the demonstrations the dolphins could freely swim around the testing tank. For Group 1 a session consisted of four to six consecutive successful demonstrations – then followed by a trial. For Group 2 a session consisted of three consecutive successful demonstrations - then followed by a trial. For Group 2, this was succeeded by two additional demonstrations, where each was followed by a trial. At the end of the session, both groups of dolphins got one more demonstration without a consecutive trial. After each demonstration the ball and the tube were taken out of the tank and the tube was re-baited out of sight of the dolphins. While re-baiting took place the dolphins in both groups were distracted by being asked to perform some training actions. During the experimental trials in phase 4, Jenny was taken to the side of the tank and received medical training (*i.e.,* lying still on her back). The trials for Group 1 were approximately three minutes long and the trials for Group 2 were approximately four minutes long. Trials normally followed demonstrations with a delay of two to five minutes for Group 1 and one to two minutes for Group 2.

If Jenny was successful in retrieving the fish during a demonstration of the target behavior, an experimenter blew a whistle (a secondary reinforcement used in dolphin training). If Jenny performed an unsuccessful demonstration, she was given the signal to repeat the task. Jenny’s performed successful demonstrations 48% of the times (23 times in Group 1 and 24 times in Group 2). Unsuccessful demonstrations were often the result of Jenny not swimming deep enough to insert the ball because she swam directly to the fish in the middle of the tube. However, whenever Jenny used the apparatus correctly - *i.e.*, whenever she took the ball and then swam to the bottom of the tube - her success rate was 100%.

While Jenny ate the fish every time she completed a successful demonstration for Group 1, in Group 2 another dolphin (Noah) stole the fish every time she performed a successful demonstration. To prevent Jenny from getting frustrated, she was manually rewarded by the experimenter after every successful demonstration. Given that in Group 2 Noah always took the fish after Jenny solved the task, it is difficult to say how much and what exactly Noah and Naomi learned from the demonstrations. To motivate the members of Group 2 to solve the task themselves without relying on Jenny, an extra 15 minute trial was conducted after the fourth demonstration session in phase 4.

#### Coding

During phase 4 (trials with demonstrations), two experimenters next to the camera loudly described which dolphins approached the apparatus and what they were doing. One of the experimenters later coded the following variables from the video recordings: in phases 1 and 3 (baseline and trial control) the experimenter coded the number of times the dolphins approached the apparatus with the ball. An approach was coded when the dolphin (with a ball) came into a radius of 1 meter from the apparatus while holding the ball in the rostrum (a rough measure of know-what and know-where). Form each approach the direction of the approach was coded: from below – the subject approaches the bottom of the tube; from the middle - the subject approached the middle of the tube where the fish was attached (a natural distractor); and from above – the subject approached the top of the tube swimming in the surface. From each approach we also coded if a ball was inserted in the tube and from where was the ball inserted (know-where 1 or 2).

Unfortunately, it was not possible to confidently code from the video recordings if the observer dolphins in each group had seen the demonstrations or not - given that dolphin’s vision field is maximized laterally.

### Results and discussion Study 1

None of the dolphins solved the task neither in the baseline nor after seeing demonstrations (in phase 4). There were no statistical differences in the individual number of approaches to the apparatus between the baseline and the trial demonstration phase (paired Wilcoxon test: *z* = 0.365, *P* = 0.875, *N* = 5). In the baseline Noah and Naomi were the only dolphins that approached the tube with the ball – thereby showing that ball approach per se does not necessitate demonstrations. In the demonstration phase four out of five dolphins approached the tube with the ball at least once – perhaps a mild social learning effect. However, the dolphins always approached the tube from above (Table 2). In the target control phase, the dolphins occasionally touched the target with their rostrum but none of them approached the tube or the target with a ball. In the extra 15 minutes trial conducted with Group 2, Naomi never approached the tube with the ball whereas Noah did so twice (from above). Thus, the members of Group 2 approached the apparatus even less often when Jenny was not present in phase 4 than during the trials where Jenny was acting as demonstrator. Therefore, we believe that the failure of the dolphins in Group 2 to solve the task was not due to their lack of motivation nor due to their reliance on Jenny to provide them with a fish.

**Table 2.** Frequency of approaches (per 60 minutes) with the ball to the apparatus during the baseline phases and demonstration phases (trials) in Study 1 (Ball up) and Study 2 (Ball down) studies. In the parentheses are the total numbers of approaches for each dolphin.

| Subject | Study 1 - Ball up |  | Study 2 - Ball down |  |
| --- | --- | --- | --- | --- |
|  | Baseline | Demo | Baseline | Demo |
| Moby | 0 | 13.3 (3) | 7.5 (4) | 5.4 (6) |
| Eva | 0 | 4.4 (1) | 0 | 0 |
| Jenny | 0 | / | 1.9 (1) | 17.2 (35) |
| Sunny | 0 | 0 | 0 | 0.7 (1) |
| Noah | 2.4 (3) | 1.6 (1) | 0 | 3.5 (6) |
| Naomi | 15.2 (19) | 3.2 (2) | 1.9 (1) | 0 |

Noah and Naomi appeared to try to solve the task in a different way than the demonstrated solution: by hitting the tube with their bodies. Their strategy did not involve using any tool and this alternative solution was used already during the baseline phase (thus, it was spontaneous). Noah and Naomi kept hitting the tube throughout the baseline phase and the trials with demonstrations. In the baseline phase they never got the fish out by hitting the tube but in the trials with demonstrations they were successful, on three occasions: twice while the tube was being put into the water and once during a trial. Unfortunately, it was not possible to count the number of hits of the tube by each dolphin given that no underwater camera was available.

## Study 2 – Ball down (human demonstrations)

### Materials and Methods

#### Subjects

Subjects were the same as in Study 1.

#### Apparatus

The apparatus was the same as in Study 1 with the exception that circles (3cm in diameter) were drawn randomly over the tube, and both ends of the tube were marked with silver tape to increase their visibility. During this second study, no target was attached to the tube. The same small basketballs (15cm in diameter) were provided as potential tools but now they were filled with saturated salt water so that they sank to the bottom of the tank (know-what 2). The intended (target) solution now involved placing these heavy balls into top of the tube (know-where 2), so that their sinking would release the fish reward. The dolphins were allowed to play and get acquainted with these balls for 10 days prior to Study 2. The balls were left in the tank on average 4 to 5 hours a day and sometimes they were left there overnight.

#### Experimental procedure

Study 2 consisted of two phases: 1.) baseline and 2.) trials with human demonstrations.

- *Baseline.* The baseline phase was conducted with the whole group and consisted of two sessions of 15 minutes each. The goal of the baseline phase was to document the actions that the dolphins displayed spontaneously toward the baited apparatus. As before, the dolphins could swim freely in and out of the adjacent performance tank. At the beginning of each session six sink balls were placed in the testing tank together with the baited apparatus.
- *Trials with human demonstrations.* This test condition included twelve sessions in which human demonstrations were provided to the dolphins. In each session dolphins were exposed to three consecutive demonstrations before participating in a trial where they were allowed to interact with the testing materials. Then they got one more demonstration followed by another trial. During the demonstrations, a human sat at the edge of the tank and released the ball under water into the tube’s upper opening, so it sank down the tube. In 30% of cases, the sinking ball did not release the fish and the demonstration was repeated. After each demonstration the ball was left in the tank. Trials were initiated one to two minutes after demonstrations ended.

In the first six sessions all six dolphins were tested together while having free access to all tanks. In these six sessions, Naomi often drew Jenny away from the tube. Since Jenny had been the subject with the most approaches to the tube with a ball (and therefore the most promising individual), we decided to take Naomi out of the experiment. However, it was not possible to separate Naomi from the rest of the group and for the next three sessions (7,8,9) Jenny and Sunny were tested alone. As Sunny seemed to present signs of distress in session 10 she was exchanged for Noah. In the last three sessions (10, 11 and 12) Jenny and Noah were separated from the other dolphins in order to watch the demonstrations. Due to these complications, the dolphins received different numbers of demonstrations: Moby, Eva and Naomi were exposed to 20 demonstrations, Sunny to 29, Noah to 30 and Jenny to 39. In the last six sessions (7,8,9,10,11 and 12) the dolphins were required to perform some training actions while the tube was being re-baited.

#### Coding

In Study 2 the same coding scheme as in Study 1 was used. In Study 2 the ball stayed in the tank after the first demonstration and so the dolphins already had the opportunity to interact with the ball during the demonstrations as well as during the trials. Thus, coding started already during the demonstrations.

### Results and discussion Study 2

Despite having ample opportunities, no dolphin solved the task with the demonstrated solution either in the baseline or the trials with demonstrations. In the baseline phase three dolphins approached the tube with the ball (all from above) and in the demonstration phase four dolphins approached the tube with the ball at least one time (Table 2). We thus did not find significant differences in the number of approaches to the tube with the ball between the baseline and the trials with demonstrations (paired Wilcoxon test: *z* = 0.405, *P* = 0.812, *N* = 6). During the trials with demonstrations, Moby, Sunny and Noah approached the tube from above, whereas Jenny approached the tube mostly from below (13 times) and from the center (21 times). On one occasion Jenny slid the ball upwards from the bottom to the top of the tube’s outside and then released the ball. Twice Jenny approached the tube from below and then tossed the ball up. Jenny also frequently hit the tube with her rostrum - both while holding the ball and sometimes without the ball. She banged the tube for the first time in the second session, but it was not until session 5 that she started hitting it regularly (more than 15 times per session). She successfully retrieved the fish one time in this manner (during a trial in session 7). Noah and Naomi successfully acquired the fish eight times with this alternative banging technique in the first six sessions (hitting the tube ca. 10 to 15 times per session).

When, later, Noah was alone with Jenny, he was never successful again. Noah and Naomi were mostly successful with the banging method (5 out of 8 times) before a trial started, that is, while the tube was being put in the water, but they almost never hit the tube during demonstrations. The occasional obtention of the fish using this alternative technique may well have affected the degree of attention paid to the demonstrations and/or reduced the need to learn a new solution for the task.

#### Comparing the results from both Ball up and Ball down studies

Dolphins’ approaches to the apparatus with the ball during both studies are presented in Table 2. The number of times that individual dolphins approached the apparatus with the ball did not differ between the two studies, neither in the baseline phase (paired Wilcoxon test: *z* = 0.365, *P* = 0.875, *N* = 6) nor in the trials with demonstration (paired Wilcoxon test: *z* = 1.1214, *P* = 0.312, *N* = 5).

## General discussion

In this study we aimed to test the abilities of six mildly enculturated dolphins to solve a tool using task either spontaneously during baselines or by socially learning the solution from or via a demonstrator (either a conspecific or a human). None of the six dolphins solved the task using demonstrated tools in the demonstrated way (using the correct know-what as a tool in the correct know-where). The failure of the dolphins in this task is unlikely to be the consequence of the tool use actions or results being too complicated for dolphins. In Study 1 all the subjects played with the air-filled balls and displayed actions with the balls very similar to those needed to retrieve the fish (e.g. submerging with the ball and letting it go again). However, the dolphins did not transfer these actions to the experimental paradigm, they failed to connect the correct know-what to the correct know-where. Given that in Study 1 the air-filled ball needed to be introduced in the tube from below in order to obtain the fish, the fact that most dolphins (except Jenny) always approached the tube from above suggests that they did not understand the task itself. Overall, across two studies we found no evidence of spontaneous tool use social learning abilities in untrained, mildly enculturated bottlenose dolphins.

The disparity between our negative results and at least some of the previous literature on dolphin social learning might however solely be the consequence of methodological difficulties (more on this below). They may also reflect the different levels of training/enculturation of the subjects included in the various studies. The test dolphins in our study were not trained to copy demonstrated actions nor received training to solve the target task in any of the two solution types, consistent with the possibility that untrained, mildly enculturated dolphins do not (or rarely) express spontaneous copying abilities in the tool domain. Experience and test-sophistication - especially in some way of know-how copying (which the dolphins we tested did not have) – could thus be important factors in eliciting know-how copying abilities in dolphins’ tool use tasks. However, this possibility is not the only one.

Another difference between our study and the previous studies reporting copying in dolphins that might have influenced our results is the time delay between the demonstrated solution and the observers’ reaction. In the present study, the dolphins were allowed to participate in the experiment one to three minutes after the demonstration (and sometimes even later). However, in previous studies (Kuczaj & Walker, 2006) the tested dolphins were allowed to interact with the testing materials immediately, i.e. while the demonstrations took place (Xitco’s, 1988). Supporting time delay as an explanation, the performance of the dolphins tested by Xitco (1988) got worse when a delay was imposed between the time the demonstrations took place and the time the dolphins could participate in the task. Future studies could systematically investigate the potentially negative effect of time delays in dolphin social learning abilities.

However, the tested dolphins in our study might simply have been unsuccessful because they sometimes saw unsuccessful demonstrations (a problem especially for Study 1), or because they did not understand the task. Horner and Whiten (2007) suggested that if apes do not have a representation of the requirements of the task (*i.e.*, if they do not understand the causality of the required actions), they then fail to perform a (demonstrated) solution to a problem. The same could be argued for the dolphins in our study. Our dolphins may not have understood the connection between the ball going through the tube and the release of the fish. Note that even the dolphin Jenny may not have fully understood the problem in the right way, since (even though she had been trained to solve the task in Study 1) she was unable to transfer this knowledge to Study 2. While this is a real possibility, note that a causal understanding of the task would have also removed the need for the dolphins to copy. Had they understood the causal demands and the causal demands alone, then they could have even independently arrived at the solutions, irrespective of demonstrations.

Another possible explanation why the dolphins did not solve the task could be that the objects the dolphins were required to use as tools in Study 1 were extensively used by the dolphins in different contexts (such as playing). Balls were not novel to the tested dolphins. As a result, it is possible that the dolphins’ performances suffered from so-called functional fixedness. Functional fixedness “occurs when the priming of a conventional / regular use for an artifact (*i.e.*, here: air-filled balls) makes it difficult to envision task-relevant atypical uses of the artifact“ (Barrett *et al*. 2008). To the dolphins, the air-filled balls’ fixed function – though perhaps achieved solely by personal experience instead of conventions 1 - was primarily to play. This may have cognitively prevented the dolphins from seeing the balls’ potential new function as a tool (see also Hanus et al. 2011). However, note that we partially accounted for this possibility, because, in Study 2 the dolphins had to solve the task using an ball object with novel, unfamiliar properties: salt-water filled balls (heavy balls; know-what 2). Since these changes had no effect in the performance of the dolphins this goes somewhat against the hypothesis that the original hindrance of the dolphins’ performance in Study 1 had been functional fixedness. Of course, even despite different fillings, the objects used in Study 1 and 2 might still have been perceptually too similar to overcome potential functional fixedness.

Last but not least, and as alluded to above, there were methodological shortcomings of our study and which could have been fully or partly responsible for our failure to find tool use social learning types in untrained, mildly enculturated dolphins. Our apparatus could be solved by the dolphins in unintended ways, and indeed this proved distractive. We hope future studies can address these shortcomings. Such studies should ensure the sturdiness of any used tool-apparatuses to prevent dolphins from being able to obtain the reward by simply hitting the apparatus. Lacking a need to use a tool in our task designed to elicit differential tool use (or at least differential know-what/know-where social learning) therefore renders our results and conclusions tentative. Motivation is a pre-requisite for meaningful tests, and here, this motivation was reduced by the possibility to access food rewards without tools – which was additionally distracting from learning the intended way(s) to solve the task.

More changes in future studies would benefit the robustness of outcomes. First, if possible, individual testing should be implemented to control the learning opportunities of each dolphin separately and to prevent the dolphins from influencing each other’s responses. Second, future studies should employ underwater cameras to capture the responses of the dolphins in detail – lacking these details curbed the conclusiveness of our findings. Third, to further investigate the effect that prior training might have on the social learning abilities of dolphins, the performance of trained and untrained dolphins should be compared (ditto for levels of enculturation). Fourth, to evaluate the effect that the type of demonstrator has on the performance of the dolphins, demonstrations from different species (conspecifics and non-conspecifics) should be provided within a single task. Fifth, age effects could be evaluated by comparing the copying abilities of both calves and adults (compare Kuczaj et al. 2012).

Once again, we do *not* hold the view that dolphins are *incapable* of spontaneous social learning. Most animals seem spontaneously capable of some social learning – it would have been instead highly surprising and unusual if dolphins had been an exception (above we showed that they are not: dolphins are clearly capable of some social learning). Yet, the fact that dolphins can learn socially in some ways does not mean that dolphins can spontaneously socially learn in all ways or that their social learning – for all transmitted types of information – always goes beyond what the individual dolphin could have discovered in their own lifetime if sufficiently motivated and in the right circumstances. This includes the special case of know-*how copying* in the tool domain. We hope that future studies will continue to investigate (using experimental controls) the tool use (and other) social learning abilities of unenculturated and untrained dolphins and their relative depth - also to better understand how tool use behaviors such as shelling or sponging may likely be transmitted in wild dolphin populations.

## Acknowledgments

We foremost thank Annette Kilian, who has sadly passed away after co-writing the first drafts of this manuscript. Due to subsequent revisions, she cannot be held to all the views expressed. We would like to express many thanks to Josep Call for his support, without whom this study would not have taken place. We thank Nathan Pyne-Carter and Alba Motes Rodrigo for improving and editing earlier manuscripts. In particular we are indebted to Tiergarten Nürnberg for letting us conduct this research and all the dolphin keepers for their help, especially to the head dolphin keeper Armin Fritz. We further thank Dr. Lorenzo von Fersen and Raik Pieszek.

## Note on this manuscript

Note that this is likely the final manuscript, as we are currently (at time of uploading) no longer intending journal submission. This is because one of the authors has sadly passed away (AK) and another author (AH) is currently no longer active in academia.

## Footnotes

1 Note that functional fixedness can be potentially established via different pathways. Fixedness may derive from purely personal experience or may be purely culturally/observationally based (note the term ”conventional“ in the quote above). In the human case, the two types could potentially mix so that individual experience may accompany cultural practices and vice versa. Here we target functional fixedness based on personal experience alone.

